# Benzaldehyde Synthases Are Encoded by Cinnamoyl-CoA Reductase Genes in Cucumber (*Cucumis sativus* L.)

**DOI:** 10.1101/2019.12.26.889071

**Authors:** Baoxiu Liu, Guo Wei, Zhongyi Hu, Guodong Wang

## Abstract

Benzaldedyde, commonly detected in plant VOC (volatile organic compounds) profiling, is derived from phenylalanine. However, the last enzymatic step for benzaldedyde formation, designated as benzaldehyde synthase, remains elusive for long time. Here, we demonstrated that cinnamoyl-CoA reductases are responsible for benzaldedyde production in cucumber (*Cucumis sativus* L.). Comprehensive tissue specificity of VOC profiling revealed that benzaldehyde was specifically accumulated in root and flower of cucumber plants. VOC-gene correlation analysis suggested that several *CCR*s are candidate genes for benzaldehyde production: *CsaCCR7* had a root-specific expression pattern while *CsaCCR9* and *CsaCCR18* showed a flower-specific expression pattern. Enzymatic assay demonstrated that CsaCCR7, CsaCCR9 and CsaCCR18 convert benzoyl-CoA to benzaldehyde. Subcellular localization experiments revealed that CsaCCR7 and CsaCCR18 are localized in cytosol, while CsaCCR9 was localized in peroxisome. In contrast to the long-standing view that CCR enzymes are involved in lignin biosynthesis in plants, it is the first time here to add a new biochemical role of CCR as benzaldehyde synthase in plants.

**Highlights:** Benzaldehyde is mainly produced in flower and root of cucumber plants.

14 genes encoding CCR enzyme from cucumber are comprehensively analyzed.

Three CsaCCRs, function as benzaldehyde synthases, utilize benzoyl-CoA as substrate to produce benzaldehyde *in vitro*.

## 1. Introduction (847 words)

Benzaldehyde (C_6_H_5_CHO, the simplest aromatic aldehyde in nature), together with other VOCs, plays a key role in plant fitness. Benzaldehyde is always detected in flower organs of many plants, and has been proved as attractants for pollinators [1, 2]. In plants, benzaldehyde biosynthesis is thought to be derived from *trans*-cinnamic acid, which was produced by degradation of phenylalanine. Currently, there are three theoretical routes to benzaldehyde formation in plants: *β*-oxidative CoA-dependent in the peroxisome (route 1), non-*β*-oxidative CoA-dependent (route 2) or CoA-independent hydratase/lyase pathways in the cytoplasm (route 3 in Fig. 1) [3]. In the petunia all functional genes involved in the peroxisomal CoA-dependent *β*-oxidation model have been identified. These included the activation of cinnamic acid catalyzed by a peroxisomal cinnamoyl-CoA ligase (CNL) [4, 5], subsequent hydration and oxidation reaction catalyzed by cinnamoyl-CoA hydratase/dehydrogenase (CHD) [6], and removal reaction catalyzed by 3-ketoacyl thiolase (KAT), followed by the production of benzoyl-CoA [7]. Moreover, thioesterases in peroxisome hydrolyzed benzoyl-CoA to benzoic acid, which might be exported via freely diffuse or unknown transporter to the cytosol [6]. However, it is not a free energy-favored reaction from benzoic acid to benzaldehyde. The cytosol CoA-dependent chain-shortening pathway (C_6_-C_3_ to C_6_-C_1_, like the *β*-oxidative pathway in peroxisome) by hydratase/lyase might another option to produce benzaldehyde in plants. This theoretic pathway is still need to be verified by experiments. Although the hydratase/lyase activity converting cinnamic acid to benzaldehyde (route 3) has been reported in *Hypericum androsaemum*, no corresponding enzyme or gene has been identified thus far [8]. Altogether, one clear fact is that benzyol-CoA existed in cytosol and peroxisome of plant cells. Whether benzyol-CoA could be directly utilized as the precursor for benzaldehyde, and the last enzymatic step (benzaldehyde synthases, BS) remain to be addressed in plants.

**Fig.1.**
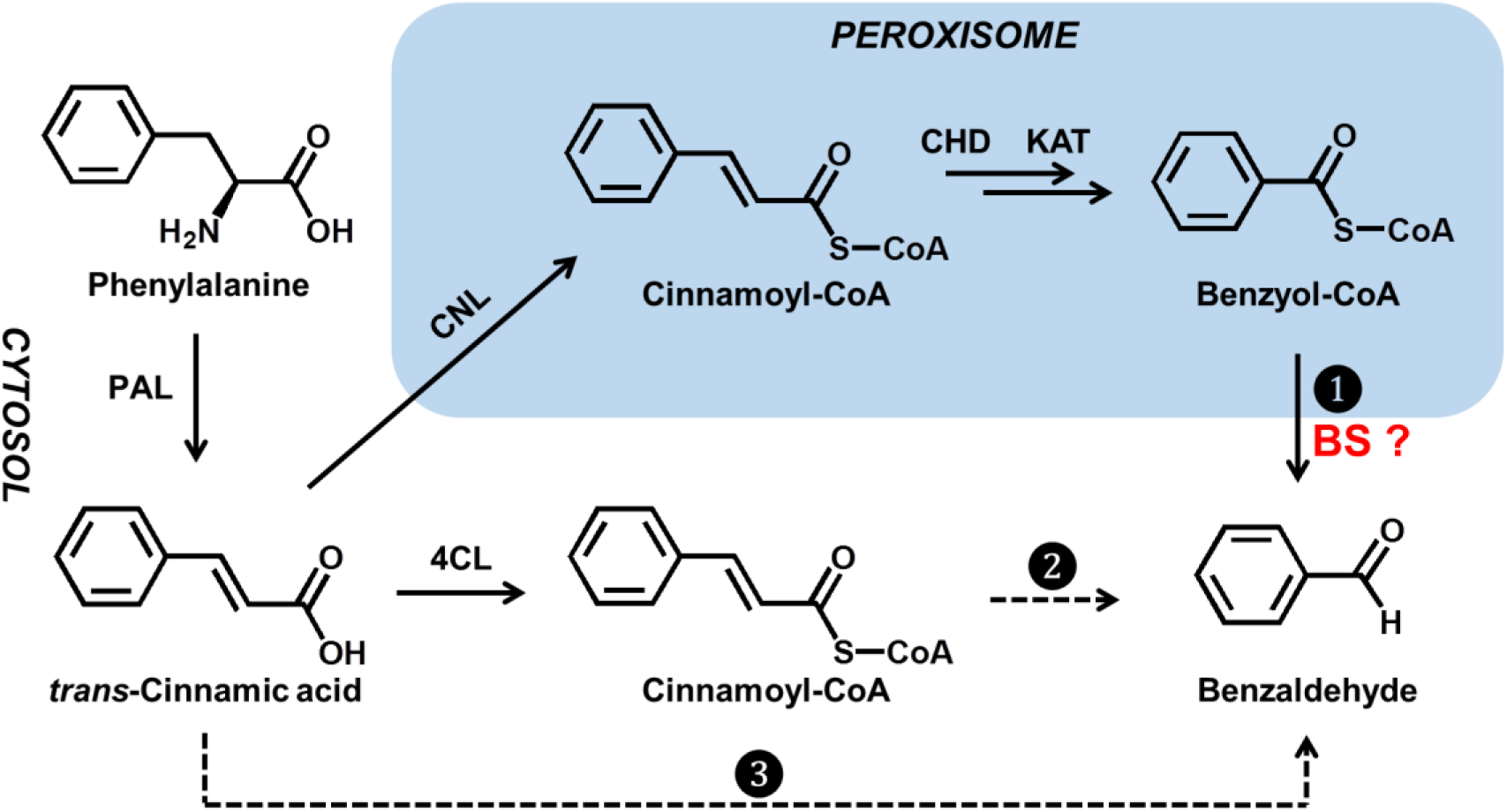
Three proposed pathways to benzaldehyde from *trans*-cinnamic acid in plants. Benzaldehyde synthase (BS), investigated in this study, was highlighted in red. Hypothetical steps are indicated by broken arrows, and multiple enzymatic steps are indicated by double arrows. CCR, cinnamoyl-CoA reductases; CHD, cinnamoyl-CoA hydratase/dehydrogenase; 4CL, 4-coumaroyl-CoA ligase; CNL, cinnamoyl-CoA ligase; KAT, 3-ketoacyl thiolase; PAL, phenylalanine ammonia lyase.

Cinnamoyl CoA reductases (CCR; EC 1.2.1.44) or CCR-like proteins, which converts hydroxycinnamoyl-CoA esters to its responding aldehydes, contributes to the first committed reaction in lignin biosynthesis [9]. CCRs in model plants (*Arabidopsis thaliana* and *Oryza staiva*) and several economic plant species (*Eucalyptus gunnii, Triticum aestivum, Pyrus bretschneideri* and *Zea mays*) have been partially identified and enzymatically analyzed [10-17]. It is noteworthy that CCR homologs form a mid-sized gene family in most plant species, for example, there are 11 CCR homologs in *Arabidopsis thaliana* [12] and 26 homologs in *Oryza sativa* [13]. Comparative and phylogenetic analyses of plant CCRs showed that CCRs could be divided into three major classes: Class one includes the *bona fide* CCR genes, while the other two classes include CCR and CCR-like, of which several genes present a high similarity to cinnamyl alcohol dehydrogenase (CAD), or dihydroflavonol reductase (DFR) [18]. To date, five common C_6_-C_3_ CoA esters, including *p*-coumaryl-CoA, caffeoyl-CoA, feruloyl-CoA, 5-hydroxyferuloyl-CoA, and sinapoyl-CoA, are always used as substrates to test CCR activities in most biochemical assays. CCR or CCR-like enzymes from the same species always showed different substrate preference. For example, AtCCR1, form *Arabidopsis thaliana*, prefer feruloyl-CoA and sinapoyl-CoA to *p*-coumaryl-CoA [12]. PtoCCR1, from *Populus tomentosa*, exhibited specific activity toward feruloyl-CoA, and no detectable activity for any other hydroxycinnamoyl-CoA esters, while PtoCCR7 showed comparable activity toward all tested hydroxycinnamoyl-CoA esters [19]. Given the similar chemical structure between C_6_-C_3_ CoA ester (cinnamoyl-CoA in Fig. 1) and benzyol-CoA and the general promiscuity of plant CCR, it is reasonable to assume plant CCR enzymes might utilize benzyol-CoA as substrate to directly produce benzaldehyde. Unfortunately, a recent study demonstrated that down-regulation of *PhCCR1* leaded to reduced amounts of lignin and phenylpropenes (C_6_-C_3_ VOCs), while no difference in benzaldehyde emission in petunia flowers (*Petunia hybrida*) [20]. However, this study did not rule out the possibility that other CCR enzymes might involve in benzaldehyde biosynthesis in petunia. With the recent release of petunia genome sequence [21], the detailed function of each member of petunia CCR family will be clarified in near future.

Cucumber (*Cucumis sativus* L.; *2n* = 14) is an important vegetable crop whose whole genome has been recently sequenced [22]. Previously, we carried out an integrative analysis of VOC profiling and transcriptome data from 23 tissues of cucumber plants to elucidate the VOC metabolic network at molecular level. We further functionally identified the genes/enzymes involved in terpenoid biosynthesis in cucumber [23]. Candidate genes responsible for other VOCs biosynthesis in cucumber remains to elucidate. In this study, we performed a genome-wide analysis of the cucumber CCR family (total 18 members) at molecular and biochemical level, and found that CsaCCR7, CsaCCR9 and CsaCCR18 function as BS by converting benzoyl-CoA to benzaldehyde. Quantitative reverse transcription-PCR and subcellular localization analysis support that both cytosol CCRs (CsaCCR7 and CsaCCR18) and peroxisomal CCR (CsaCCR9) are probably the main contributor for benzaldehyde production in different tissues of cucumber. Our data not only demonstrate that the CCR probably participate in benzyaldehyde biosynthesis in cucumber, but also provide novel insight into the whole benzenoid network in plants.

## 2. Materials and methods (714 words)

### 2.1 Plant Materials and Chemicals

In this study, the growth of cucumber plants (*Cucumis sativus* ‘9930’) and *Arabidopsis thaliana* (Col-0 ecotype, and Cs16259 (peroxisome marker line)) for CsaCCR7, CsaCCR9, and CsaCCR18 transient transformation were carried out as described previously [23, 24]. All commercial available chemicals used in this study were purchased from Sigma-Aldrich. Plant sample collection, VOC sampling and GC-MS analysis were also performed as described previously [23]. Briefly, 100 ± 0.5 mg tissue powders was weighted into a 4 mL glass vial and filled with 400 μL 20% NaCl solution (2-heptone was added as an internal standard; final concentration 125 ng μL^-1^). A cleaned fiber was inserted into the glass vial (preheated at 30°C for 5 min) and exposed to the headspace at 30°C for 30 min. The fiber was then introduced into the injector port of a GC/MS instrument (Agilent 7890A GC-5975) and held for 30 seconds. The GC-MS data acquisition and analysis were peformed as previously described [23].

### 2.2 Gene isolation, Sequence Alignment and Phylogenetic Analysis

Isolation of RNA and cDNA synthesis were carried out as previously described [25]. Genome-wide screening of CCR and its homologous have been performed based on cucumber genome [22]. To obtain the full-length sequences of CCR from cucumber, the ORF of CCR obtained by RT-PCR were cloned into pEASY-Blunt vector (TransGen Biotech) and verified by sequencing of at least five independent clones (primer information see Table S2).

Deduced protein sequences of OsCCRs and functional CCRs identified from other plant species were retrieved from the MSU RGAP database (http://rice.plantbiology.msu.edu/) and the National Center for Biotechnological Information (https://www.ncbi.nlm.nih.gov/) database, respectively. Multiple amino acid sequence alignment was performed with Clustal-W, and a phylogenetic analysis was conducted with MEGA ver. 6.0 using the neighbor-joining method [26].

All CsaCCR sequences reposted in this study have been deposited in the GenBank database (accession nos. MN868259 – MN868276).

### 2.3 Quantitative RT-PCR Analysis

Real-time PCR analyses were performed using Ultra SYBR Mixture (CWBio) on a CFX96 Real-Time PCR Detection System (BioRad) following the manufacturer’s instructions. Ct values were calculated using the Bio-Rad real-time analysis software. Comparative Ct method was used for relative gene expression analysis by normalizing to the cucumber *UEP* (ubiquitin extension protein) gene (GenBank accession No. AY372537). Every measurement was performed with at least three biological replicates. Primers are listed in Table S2.

### 2.4 Subcellular localization

The ORF of the GFP (green fluorescent protein) gene was fused to the C-terminal of the *CsaCCR7* and *CsaCCR18*, under control of the CaMV 35S promoter (pJIT163-hGFP vector). The ORF of CsaCCR9 was fused to mCherry (pJIT163-mCherry vector) at the C-terminal. Arabidopsis leaf protoplast preparation, transformation, and image assay using laser scanning confocal microscopy were performed as described previously [27]. Briefly, mesophyll protoplasts freshly isolated from rosette leaves of 4-week-old Arabidopsis plants. The fresh prepared protoplasts are transfected with 10 μg plasmid using a PEG-calcium–mediated transfection method. Living cellular image of GFP fusion proteins are observed under Axio Imager Z2 fluorescence microscopy (Zeiss). Localization was determined by surveying more than 50 protoplasts. Primers are listed in Table S2.

### 2.5 *In Vitro* CCR Enzyme Assays

The *E.coli* (BL21 DE3) transformants harboring the pEasy-E1-CsaCCRs constructs (full length CsaCCRs fused with a His-tag at *N*-terminal) were grown at 37°C until an OD_600_ of 0.4-0.6 in LB medium containing ampicillin (50 µg/mL) was achieved. Isopropyl β-D-thiogalactopyranoside (IPTG, 0.1 mM) was then added in the culture for protein induction. After additional incubation at 16°C for 16 h, the cells were harvested by centrifugation (4°C, 5,000 *g* for 15 min). Cell pellets were resuspended in phosphatebuffered saline (137 mM NaCl, 2.7 mM KCl, 10 mM Na_2_HPO_4_, 2 mM KH_2_PO_4_). The resuspended cells were sonicated on ice, and the crude protein extracts were obtained by centrifugation (4°C, 15,900 *g* for 20 min). The crude protein samples were used for *in vitro* enzyme assays and the non-induced E. coli extract was used as a control. For kinetic analysis, proteins were purified. The crude protein samples were mixed with Ni-NTA agarose beads and purified as described previously [23].

CsaCCR activity was carried out according to the methods described by Lüderitz and Grisebach, using benzoyl-CoA as substrates [28]. The reaction mixture consisted of 0.1 mM NADPH, 30 µM benzoyl-CoA, and 5 µg of crude CsaCCR proteins in 100 mM sodium/potassium phosphate buffer (pH 6.25) to a total volume of 100 µL. The enzyme reactions were carried out at 30 °C for 30 min. Benzaldehyde was analyzed using GC/MS. For the determination of apparent *K*_*m*_ values, the reaction was initiated by an addition of benzoyl-CoA, and decreases in OD_340_ were monitored for 10 min. The substrates were used at concentrations of 5–50 µM. *Km* and Vmax were determined by extrapolation from Lineweaver-Burk plots. The enzyme assays were carried out in quadruplicate and the result represented the mean ± standard deviation.

## 3. Results and Discussion

### 3.1 Benzaldehyde is abundant in cucumber roots and flowers

We previously used SPME-GC-MS to analyze the different tissues and development time of cucumber. The distribution of volatile compounds in different tissues was also calculated. The amount 85 volatile compounds, mostly in the form of aldehyde or alcohol, were detected in 23 tissues [23]. Some volatile compounds were specifically accumulated in some tissues, such as nonaldehyde compounds are mainly accumulated in fruits, while hexanal compounds are mainly accumulated in leaves (Fig. 2A). GC-MS analysis showed that benzaldehyde and benzyl alcohol (probably derived from benzaldehyde by an oxidoreductase) are accumulated in roots and male/female flowers. The similar pattern of benzaldehyde and benzyl alcohol also suggested that the both chemicals’ biosynthetic pathways are regulated by a same system. The tissue specificity of benzaldehyde suggests the genes involved in benzaldehyde biosynthetic pathway should have a root-predominant and/or flower predominant, especially for those downstream structural genes.

**Fig. 2.**
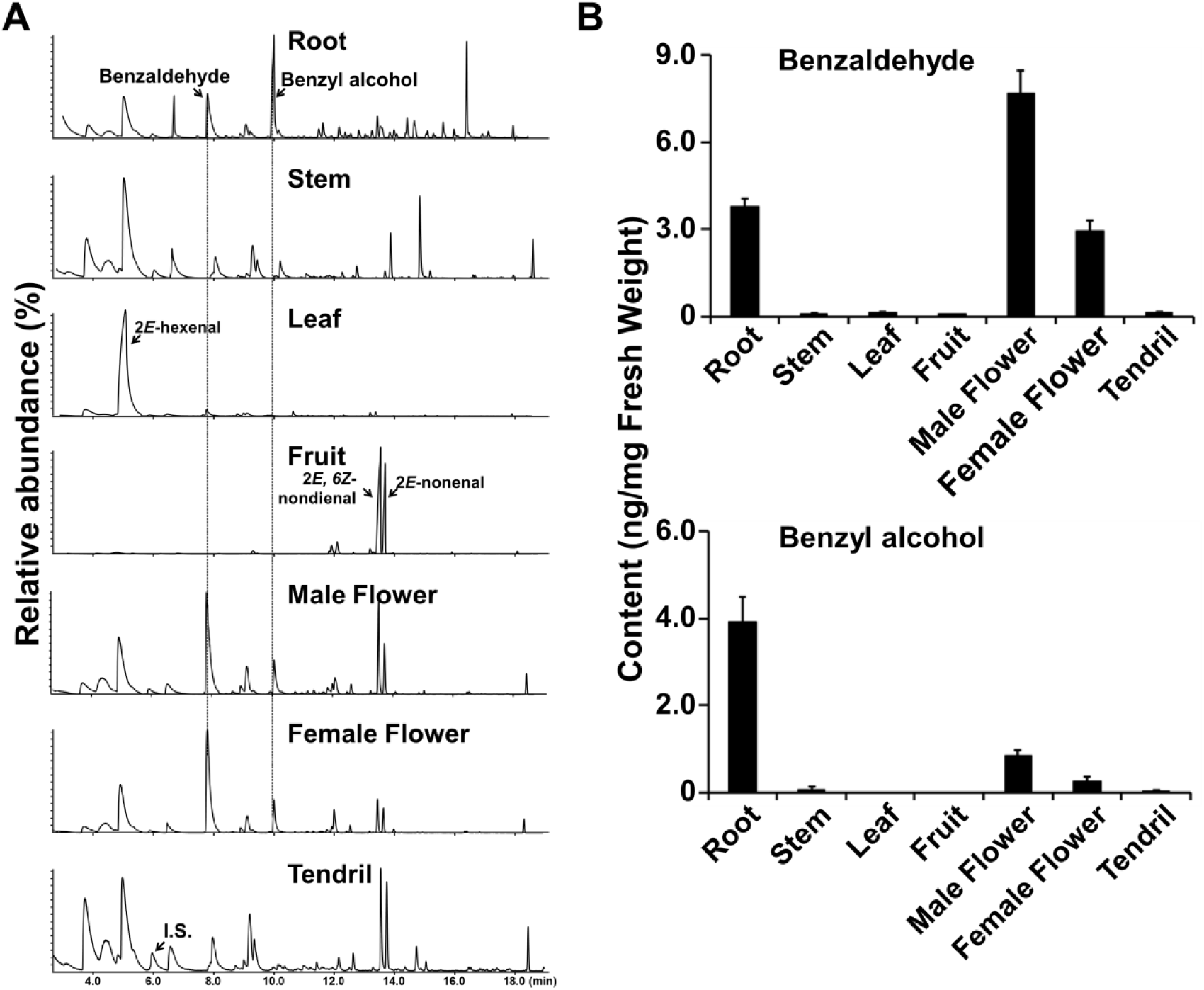
Endogenous benzaldehyde measurements in seven representative cucumber tissues. (A) The VOCs profiling in seven representative tissues (root, stem, leaf, fruit (20 day-after-pollination), male flower, female flower and tendrill) of cucumber variety 9930 using SPME-GC-MS (TIC mode), the position of several well-known VOCs are marked with arrows. I.S., internal standard (2-heptone). The highest peak in each chromatography was set as 100%. (B) Quantitative measurement of benzaldehyde (upper panel) and benzoyl alcohol (lower panel) in seven tissues of cucumber variety 9930, data are presented as mean ± S.D. (*n* = 3).

### 3.2 Cucumber CCR genes cloning and analysis

Previous studies in petunia flowers showed that benzaldehyde content in *PhCHD*-RNAi and *PhKAT1*-RNAi (both PhCHD and PhKAT1 are localized in peroxisome and involved in *β*-oxidation process) transgenic plants were significantly decreased [6, 7, 29]. As shown in Fig. 1, benzaldehyde is most likely derived from the CoA-dependent *β*-oxidation process of cinnamic acid in plant cells (the route 1 in Fig. 1), and the last step from benzoyl-CoA to benzaldehyde might be catalyzed by a CCR enzyme. To test this hypothesis, we searched the cucumber genome, and total 18 CCR-encoding genes were found (detailed information see Table S2). The expression patterns of these 18 *CsaCCR* genes in seven tested cucumber tissues were extracted from our previous study [23], and no transcripts of *CsaCCR10* and *CsaCCR11* could be detected in seven tested tissues (Fig. S1). The result showed that at least seven *CsaCCR*s (*CsaCCR1, CsaCCR6-9, CsaCCR13, CsaCCR18*) showing root-predominant and/or flower-predominant pattern were determined as benzaldehyde biosynthesis candidate genes for next study. It is noteworthy that *CsaCCR6, CsaCCR10*, and *CsaCCR15* were excluded from further study due to the too short protein sequences they encoded (Table S2). Subcellular prediction by TargetP (www.cbs.dtu.dk/services/TargetP) and WolF PSORT (http://wolfpsort.org/) suggested most CsaCCR proteins are cytosol proteins, except that there is an *N*-terminal signal peptide for several CsaCCRs, including CsaCCR3, 7-9, 12, and 13 (Table S2).

### 3.3 CsaCCR7, CsaCCR9 and CsaCCR18 function as benzaldehyde synthase (BS) *in vitro*

Phylogenetic analysis shows that CsaCCR7, CsaCCR9 and CsaCCR18 are grouped with all functional identified plant CCRs, which utilize hydroxycinnamoyl-CoAs as substrate (Fig. 3A). This result also hints that CsaCCR7, CsaCCR9 and CsaCCR18 might involve in lignin biosynthesis in cucumber. To test the hypothesis whether one or more CsaCCRs involve in benzlaldehyde biosynthesis, we further checked the 14 CCR activity (using the crude protein for large-scale activity screening) *in vitro* using benzoyl-CoA as substrate, and discovered that CsaCCR7, CsaCCR9 and CsaCCR18 catalyzed benzoyl-CoA to produce benzaldehyde (Fig. 3B and 3C). The biochemical constants of three enzymes were further determined using benzoyl-CoA as substrate, and the results showed that the catalytic effenciency (*K*_*cat*_/*K*_*m*_) of CsaCCR18 was two folds higher than that of CsaCCR7 and CsaCCR9 (Table 1). It is noteworthy that catalytic effenciency of these three CsaCCRs was similar to the CCRs from other plant species when testing with hydroxylcinnamoyl-CoAs [13]. Based on these biochemical data, we tentatively re-designated CsaCCR7, CsaCCR9 and CsaCCR18 as CsaBS1 (benzaldeheyde synthase), CsaBS2, CsaBS3, respectively. Definitely, this conclusion merits further investigation at genetic level *in planta*.

**Table 1.**
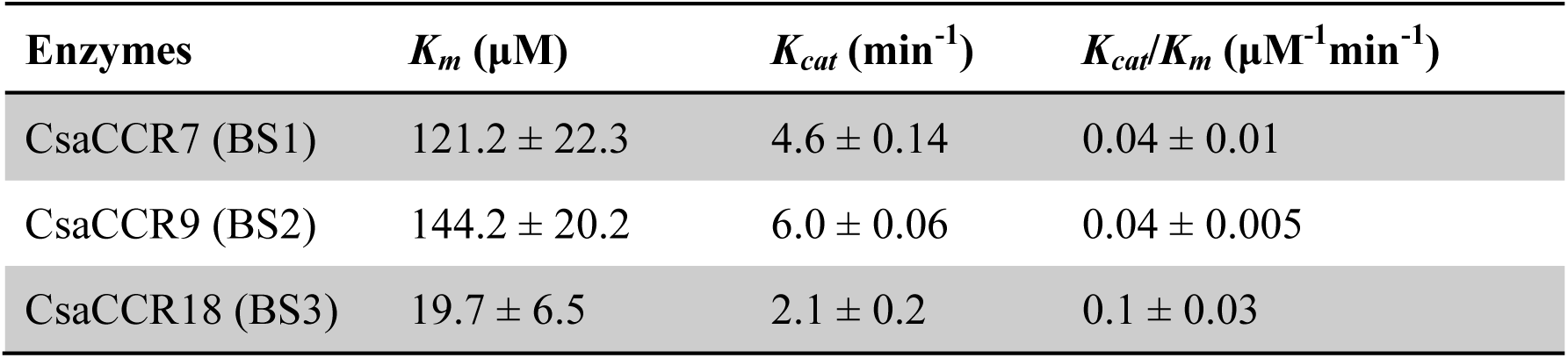
Kinetic parameters for three cucumber CCRs with benzoyl-CoA ester. Purified CsaCCR proteins were used in this assay (Fig. S3). Data are presented as means ± SD of triplicate experiments.

**Fig.3.**
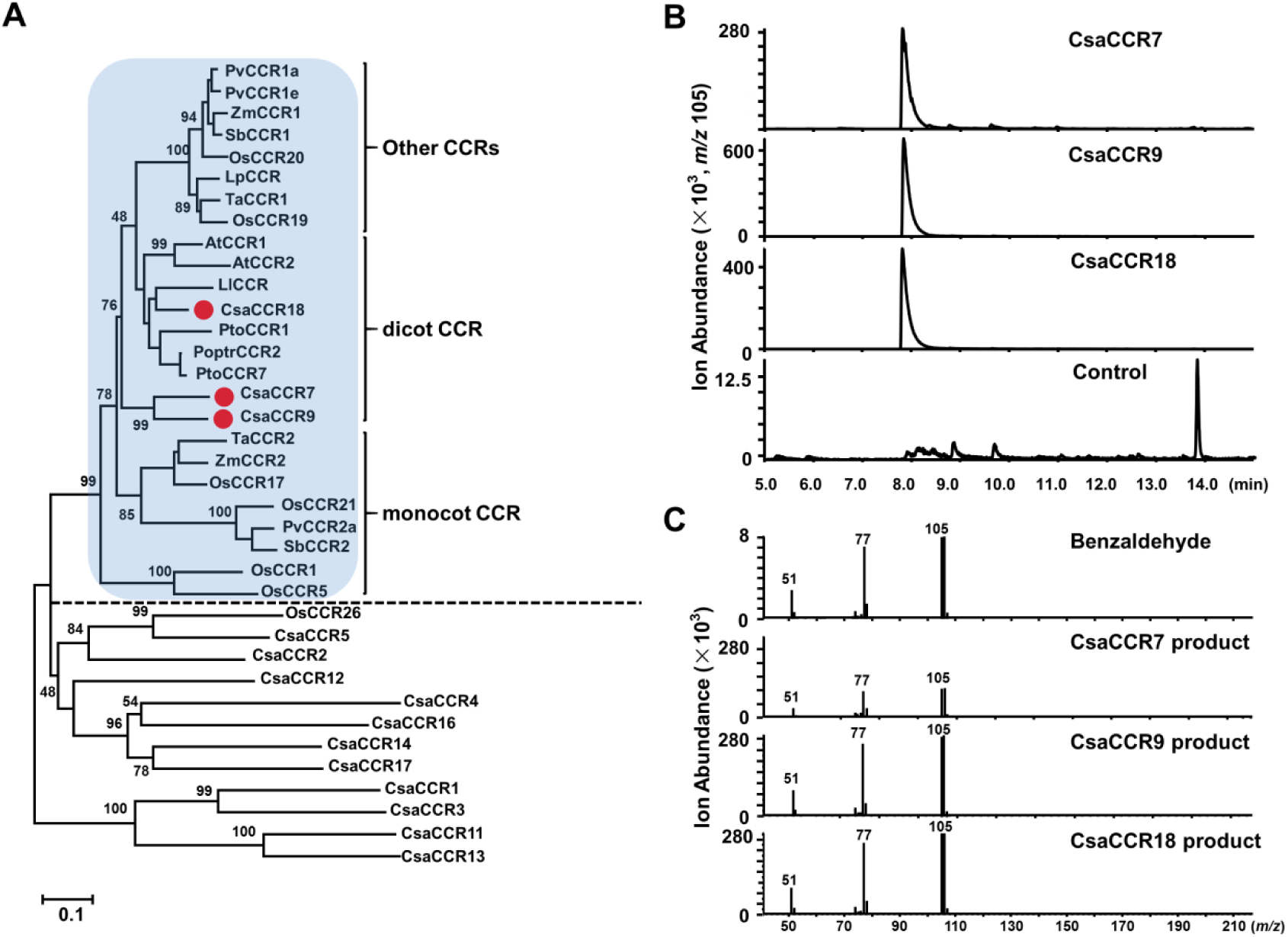
Characterization of CsaCCR7, CsaCCR9 and CsaCCR18. (A) Phylogenetic analysis of CsaCCRs and functional identified CCR proteins from other plant species. The neighbor-joining tree was built using MEGA6. Bootstrap values (based on 10,000 replicates) are shown for corresponding nodes. Class one CCRs are indicated with blue shade, and CsaCCR7, CsaCCR9 and CsaCCR18 are highlighted with red dots. Abbreviations: At, *Arabidopsis thaliana*; Ll, *Leucaena leucocephala;* Os, *Oryza sativa*; Poptr, *Populus trichocarpa*; Pto, *Populus tomentosa*; Pv, *Panicum virgatum*; Sb, *Sorghum bicolor*; Ta, *Triticum aestivum*; Zm, *Zea mays.* (B), (C) CsaCCR7, CsaCCR9 and CsaCCR18 convert benzoyl-CoA to benzaldehyde, chromatogram of selected ions of *m/z* 105, and verified by mass-spectrum comparison with reference chemical. Crude CsaCCR proteins were used in this assay, and one cucumber gene (*Csa3G611340.1*, encoding phenylacetaldehyde synthase) unrelated to benzenoid biosynthesis was used as control. This experiment was repeated at least three times with similar results.

### 3.4 Molecular characterization of three CsaBSs

Quantitative RT-PCR analysis firstly confirmed that *CsaBS1, CsaBS2*, and *CsaBS3* show root-predominant, male flower-predominant, and female-predominant pattern, which suggested all three *CsaBS*s might be responsible for benzyaldehyde production in different tissues (Fig. 4A). It is well-known that CCR is a key enzyme involved in lignin synthesis and always located in cytoplasm. However, benzoyl-CoA produced by cinnamic acid through the process of *β*-oxidation stays in the peroxisome. To test whether these above-mentioned BSs exist in a unique subcellular compartment (in peroxisome or others), we then constructed subcellular localizaiton vectors for these three *BS* genes, and transformed the protoplasts of wild-type *Arabidopsis thaliana* or Cs16259 line (a plant with stable expression of peroxisome marker gene). The results revealed that CsaBS2 protein was located in the peroxisome of the cell, while CsaBS1 and CsaBS3 were located in the cytoplasm, as predicted.

**Fig. 4.**
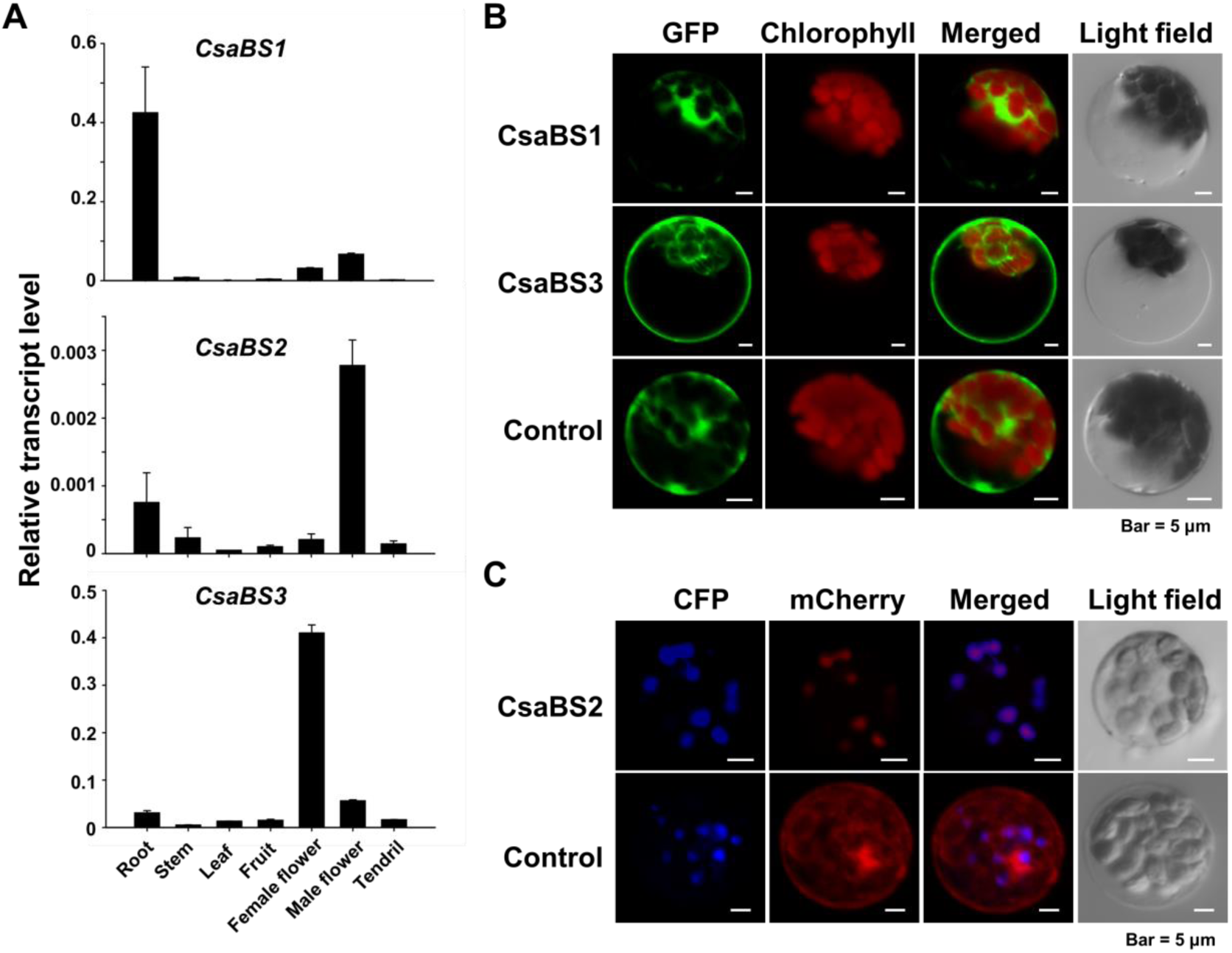
Characterization of CsaBS1, CsaBS2 and CsaBS3. (A) Quantitative RT-PCR analysis of three cucumber *BS* genes in different tissues of cucumber plants. Transcript levels are expressed relative to cucumber *UEP* (ubiquitin extension protein) transcripts (mean ± S.D., *n* = 3). (B) Subcellular localization of CsaBS1 and CsBS3 in Arabidopsis leaf mesophyll protoplasts as revealed by laser confocal microscopy. Chloroplasts are revealed by red chlorophyll auto fluorescence, and free GFP was used as a control. (C) Subcellular localization of CsaBS2 in Arabidopsis leaf mesophyll protoplasts as revealed by laser confocal microscopy. Peroxisomes are revealed by peroxisome marker protein, free CFP (cyan fluorescent protein) was used as a control.

Given the general promiscuity of plant CCRs, the substrate availability will be a key factor for the physiological functions of CCR enzymes *in planta*. In plant cells, the evidence generated from the chemical analysis of transgenic plants supported that benzoyl-CoA was produced in peroxisomes via CoA dependent β-oxidation process [2, 7]. Although the active transporter for transportation of benzoyl-CoA from peroxisome to cytosol remains unclear, the occurrence of benzoyl-CoA in both cellular compartments (peroxisome and cytosol) was demonstrated. Here we demonstrated two types of BSs (interestedly, one in peroxisome, and another in cytosol) are responsible for benzaldehyde in cucumber. Our study also provide novel insight to the unsolved vanillin (one aromatic aldehyde with similar structure to benzaldehyde) biosynthesis [30].

## Author contributions

G.Wa. designed the study; B.L., G.We., and Z.H. performed the research; B.L., and G.Wa. analyzed the data; B.L., and G.Ws. wrote the paper with constructive input from all authors.

## Declaration of competing interest

The authors declare that they have no conflict of interest.

## Acknowledgement

This work was financially supported by Key R&D Program of Shandong Province (grant No. 2019JZZY020610), National Key R&D Program of China (grant No. 2018YFA0900600), and the State Key Laboratory of Plant Genomics of China (Grants No. SKLPG2016A-13 and SKLPG2016B-13) to G.Wa.

## Supplementary data

Fig. S1. Tissue specificity of *CsaCCR* genes.

Fig. S2. *In vitro* CCR assays with Benzyol-CoA as substrate.

Fig. S3. Gel analysis of purified recombinant CsaCCR7, 9 and 18. Table S1. Primers used in this study.

Table S2. CsaCCRs subcellular localization prediction by using Traget P and Wolf PSORT software.

## References

[1] E. Pichersky, J.P. Noel, N. Dudareva, Biosynthesis of plant volatiles: Nature’s diversity and ingenuity, Science, 311 (2006) 808–811.

[2] N. Dudareva, A. Klempien, J.K. Muhlemann, I. Kaplan, Biosynthesis, function and metabolic engineering of plant volatile organic compounds, New Phytologist, 198 (2013) 16–32.

[3] J.R. Widhalm, N. Dudareva, A Familiar Ring to It: Biosynthesis of Plant Benzoic Acids, Molecular Plant, 8 (2015) 83–97.

[4] T.A. Colquhoun, D.M. Marciniak, A.E. Wedde, J.Y. Kim, M.L. Schwieterman, L.A. Levin, A. Van Moerkercke, R.C. Schuurink, D.G. Clark, A peroxisomally localized acyl-activating enzyme is required for volatile benzenoid formation in a Petuniahybrida cv. oMitchell Diploid’flower, J. Exp. Bot., 63 (2012) 4821–4833.

[5] A. Klempien, Y. Kaminaga, A. Qualley, D.A. Nagegowda, J.R. Widhalm, I. Orlova, A.K. Shasany, G. Taguchi, C.M. Kish, B.R. Cooper, J.C. D’Auria, D. Rhodes, E. Pichersky, N. Dudareva, Contribution of CoA Ligases to Benzenoid Biosynthesis in Petunia Flowers, Plant Cell, 24 (2012) 2015–2030.

[6] A.V. Qualley, J.R. Widhalm, F. Adebesin, C.M. Kish, N. Dudareva, Completion of the core beta-oxidative pathway of benzoic acid biosynthesis in plants, Proc. Natl. Acad. Sci. USA, 109 (2012) 16383–16388.

[7] A. Van Moerkercke, I. Schauvinhold, E. Pichersky, M.A. Haring, R.C. Schuurink, A plant thiolase involved in benzoic acid biosynthesis and volatile benzenoid production, Plant J., 60 (2009) 292–302.

[8] A.M.A. Abd El-Mawla, L. Beerhues, Benzoic acid biosynthesis in cell cultures of Hypericum androsaemum, Planta, 214 (2002) 727–733.

[9] E. Lacombe, S. Hawkins, J. VanDoorsselaere, J. Piquemal, D. Goffner, O. Poeydomenge, A.M. Boudet, J. GrimaPettenati, Cinnamoyl CoA reductase, the first committed enzyme of the lignin branch biosynthetic pathway: Cloning, expression and phylogenetic relationships, Plant J., 11 (1997) 429–441.

[10] L.G. Li, X.F. Cheng, S.F. Lu, T. Nakatsubo, T. Umezawa, V.L. Chiang, Clarification of cinnamoyl co-enzyme a reductase catalysis in monolignol biosynthesis of aspen, Plant and Cell Physiology, 46 (2005) 1073–1082.

[11] T. Kawasaki, H. Koita, T. Nakatsubo, K. Hasegawa, K. Wakabayashi, H. Takahashi, K. Urnemura, T. Urnezawa, K. Shimamoto, Cinnamoyl-CoA reductase, a key enzyme in lignin biosynthesis, is an effector of small GTPase Rac in defense signaling in rice, Proc. Natl. Acad. Sci. USA, 103 (2006) 230–235.

[12] M.A. Costa, R.E. Collins, A.M. Anterola, F.C. Cochrane, L.B. Davin, N.G. Lewis, An in silico assessment of gene function and organization of the phenylpropanoid pathway metabolic networks in Arabidopsis thaliana and limitations thereof, Phytochemistry, 64 (2003) 1097–1112.

[13] H.L. Park, S.H. Bhoo, M. Kwon, S.W. Lee, M.H. Cho, Biochemical and Expression Analyses of the Rice Cinnamoyl-CoA Reductase Gene Family, Front Plant Sci, 8 (2017).

[14] F.S. Poke, D.P. Martin, D.A. Steane, R.E. Vaillancourt, J.B. Reid, The impact of intragenic recombination on phylogenetic reconstruction at the sectional level in Eucalyptus when using a single copy nuclear gene (cinnamoyl CoA reductase), Mol Phylogenet Evol, 39 (2006) 160–170.

[15] Q.H. Ma, Characterization of a cinnamoyl-CoA reductase that is associated with stem development in wheat, J. Exp. Bot., 58 (2007) 2011–2021.

[16] X.Q. Su, Y. Zhao, H. Wang, G.H. Li, X. Cheng, Q. Jin, Y.P. Cai, Transcriptomic analysis of early fruit development in Chinese white pear (Pyrus bretschneideri Rehd.) and functional identification of PbCCR1 in lignin biosynthesis, Bmc Plant Biology, 19 (2019).

[17] J.R. Andersen, I. Zein, G. Wenzel, B. Darnhofer, J. Eder, M. Ouzunova, T. Lubberstedt, Characterization of phenylpropanoid pathway genes within European maize (Zea mays L.) inbreds, Bmc Plant Biology, 8 (2008).

[18] A. Barakat, N.B.M. Yassin, J.S. Park, A. Choi, J. Herr, J.E. Carlson, Comparative and phylogenomic analyses of cinnamoyl-CoA reductase and cinnamoyl-CoA-reductase-like gene family in land plants, Plant Sci., 181 (2011) 249–257.

[19] N. Chao, N. Li, Q. Qi, S. Li, T. Lv, X.N. Jiang, Y. Gai, Characterization of the cinnamoyl-CoA reductase (CCR) gene family in Populus tomentosa reveals the enzymatic active sites and evolution of CCR, Planta, 245 (2017) 61–75.

[20] J.K. Muhlemann, B.D. Woodworth, J.A. Morgan, N. Dudareva, The monolignol pathway contributes to the biosynthesis of volatile phenylpropenes in flowers, New Phytologist, 204 (2014) 661–670.

[21] A. Bombarely, M. Moser, A. Amrad, M. Bao, L. Bapaume, C.S. Barry, M. Bliek, M.R. Boersma, L. Borghi, R. Bruggmann, M. Bucher, N. D’Agostino, K. Davies, U. Druege, N. Dudareva, M. Egea-Cortines, M. Delledonne, N. Fernandez-Pozo, P. Franken, L. Grandont, J.S. Heslop-Harrison, J. Hintzsche, M. Johns, R. Koes, X.D. Lv, E. Lyons, D. Malla, E. Martinoia, N.S. Mattson, P. Morel, L.A. Mueller, J. Muhlemann, E. Nouri, V. Passeri, M. Pezzotti, Q.Z. Qi, D. Reinhardt, M. Rich, K.R. Richert-Poggeler, T.P. Robbins, M.C. Schatz, M.E. Schranz, R.C. Schuurink, T. Schwarzacher, K. Spelt, H.B. Tang, S.L. Urbanus, M. Vandenbussche, K. Vijverberg, G.H. Villarino, R.M. Warner, J. Weiss, Z. Yue, J. Zethof, F. Quattrocchio, T.L. Sims, C. Kuhlemeier, Insight into the evolution of the Solanaceae from the parental genomes of Petunia hybrida, Nature Plants, 2 (2016).

[22] S. Huang, R. Li, Z. Zhang, L. Li, X. Gu, W. Fan, W.J. Lucas, X. Wang, B. Xie, P. Ni, The genome of the cucumber, Cucumis sativus L, Nat Genet, 41 (2009) 1275–1281.

[23] G. Wei, P. Tian, F. Zhang, H. Qin, H. Miao, Q. Chen, Z. Hu, L. Cao, M. Wang, X. Gu, S. Huang, M. Chen, G. Wang, Integrative analyses of nontargeted volatile profiling and transcriptome data provide molecular insight into VOC diversity in cucumber plants (*Cucumis sativus*), Plant Physiol., 172 (2016) 603–618.

[24] B.K. Nelson, X. Cai, A. Nebenfuhr, A multicolored set of in vivo organelle markers for co-localization studies in Arabidopsis and other plants, Plant J., 51 (2007) 1126–1136.

[25] G. Wang, L. Tian, N. Aziz, P. Broun, X. Dai, J. He, A. King, P.X. Zhao, R.A. Dixon, Terpene biosynthesis in glandular trichomes of hop, Plant Physiol., 148 (2008) 1254–1266.

[26] K. Tamura, G. Stecher, D. Peterson, A. Filipski, S. Kumar, MEGA6: Molecular Evolutionary Genetics Analysis Version 6.0, Molecular Biology and Evolution, 30 (2013) 2725–2729.

[27] S.D. Yoo, Y.H. Cho, J. Sheen, Arabidopsis mesophyll protoplasts: a versatile cell system for transient gene expression analysis, Nature Protocols, 2 (2007) 1565–1572.

[28] T. Luderitz, H. Grisebach, Enzymic-Synthesis of Lignin Precursors Comparison of Cinnamoyl-Coa Reductase and Cinnamyl Alcohol - Nadp+ Dehydrogenase from Spruce (Picea-Abies L) and Soybean (Glycine-Max L), Eur. J. Biochem., 119 (1981) 115–124.

[29] I. Orlova, A. Marshall-Colon, J. Schnepp, B. Wood, M. Varbanova, E. Fridman, J.J. Blakeslee, W.A. Peer, A.S. Murphy, D. Rhodes, E. Pichersky, N. Dudareva, Reduction of benzenoid synthesis in petunia flowers reveals multiple pathways to benzoic acid and enhancement in auxin transport, Plant Cell, 18 (2006) 3458–3475.

[30] A. Kundu, Vanillin biosynthetic pathways in plants, Planta, 245 (2017) 1069–1078.

