## supplemental data for "Benzaldehyde Synthases Are Encoded by Cinnamoyl-CoA Reductase Genes in Cucumber (*Cucumis sativus* L.)"

**Supplementary Information**

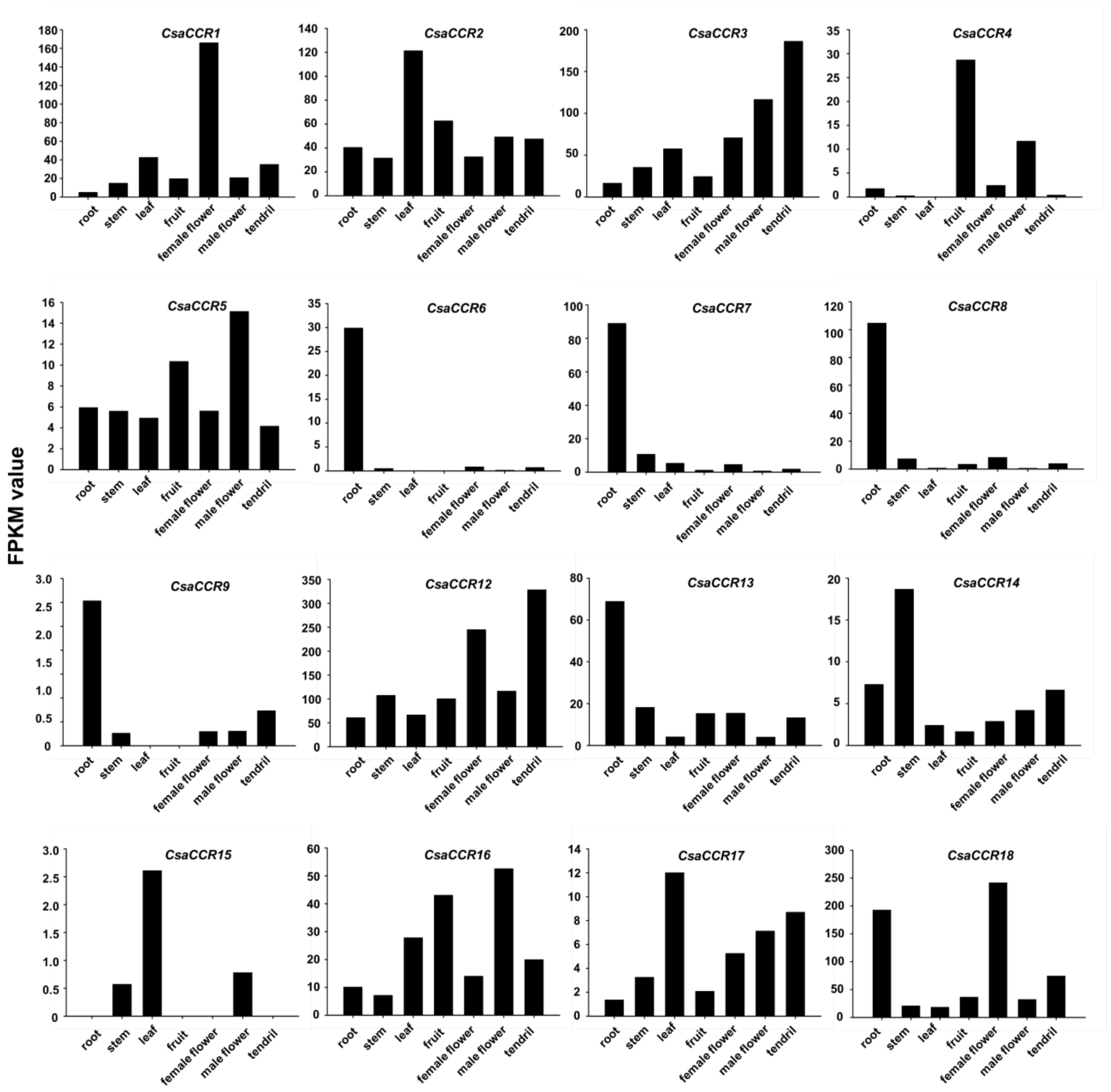


**Fig. S1. Tissue specificity of *CsaCCR* genes.** The data was extracted from RNAseq dataset generated by Wei et al., (2016). It is noteworthy that no *CsaCCR10* and *CsaCCR11* transcripts were detected in this experiment. FPKM, fragments per kilobase of transcript per million fragments mapped.

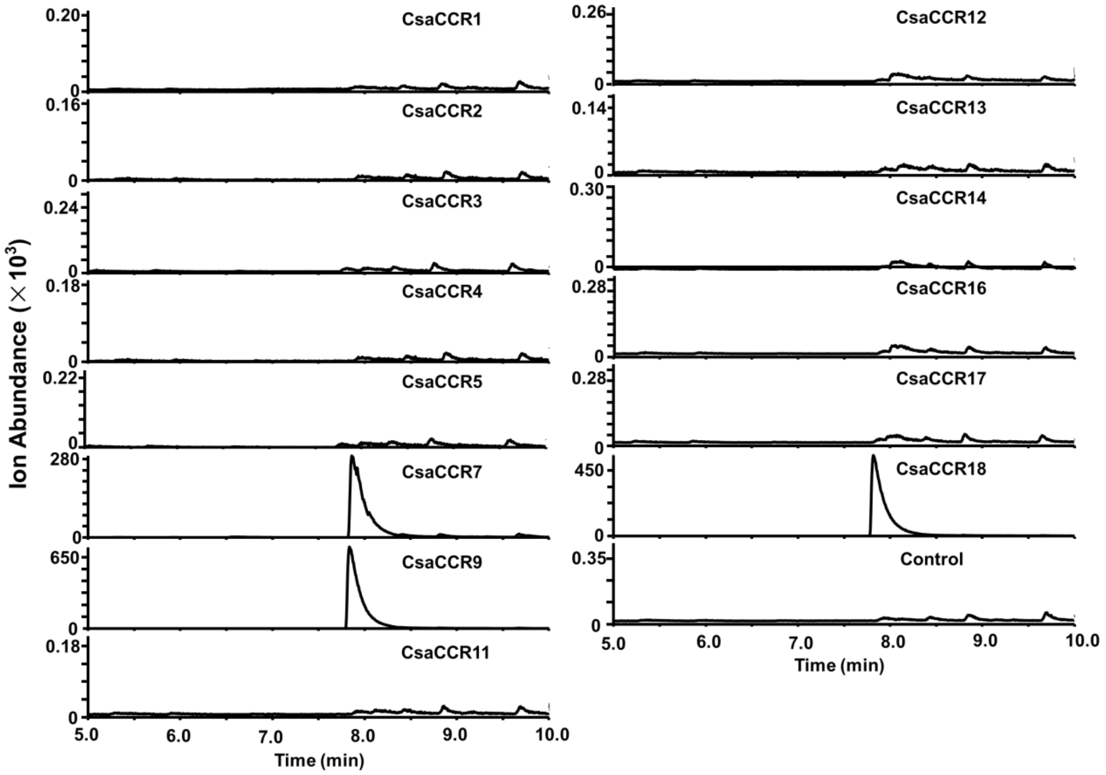


**Fig. S2. *In vitro* CCR assays with Benzyol-CoA as substrate.** Crude CsaCCR proteins were used in this assay. The enzyme product, benzaldehyde was analyzed with GC-MS with SIM mode (*m/z* 105).

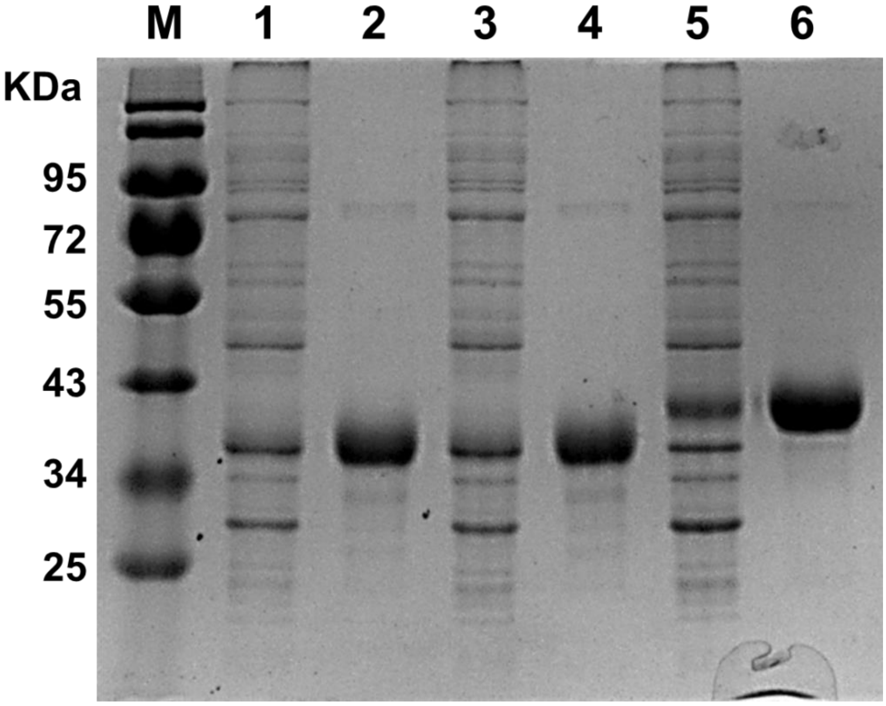


**Fig. S3. Gel analysis of purified recombinant CsaCCR7, 9 and 18.** The *N*-terminal His-tagged *CsaCCR7*, *CsaCCR9* and *CsaCCR18* were expressed in *E. coli*. The recombinant CCR proteins were purified by $Ni^{2+}$-affinity chromatography. M, Protein marker; 1, CsaCCR7 total protein; 2, purified CsaCCR7; 3, CsaCCR9 total protein ; 4, purified CsaCCR9; 5, CsaCCR18 total protein ; 6, purified CsaCCR18.

**Table S1. Primers used in this study.**

| Primer name | Sequence (5’ to 3’) |
|---|---|
| **For CCR gene cloning (into pEASY vector)** | |
| CsaCCR1-FL-For | ATGGATCATTCAAAGCCAACTCTTTG |
| CsaCCR1-FL-Rev | TTAAGCAGTGCCTTCGATGAGTTTGTG |
| CsaCCR2-FL-For | ATGTCCACCACCCAGAATCCTGAAC |
| CsaCCR2-FL-Rev | TCAGGAAATATATCCCTTCTTTTTAGATCC |
| CsaCCR3-FL-For | ATGGCTCCTCCCGCTAAGACCG |
| CsaCCR3-FL-Rev | TCAACATGAATCTCCACTTTCAAAATCC |
| CsaCCR4-FL-For | ATGGAAGCAGAAGAAGAAGATGAAGAAAAT |
| CsaCCR4-FL-Rev | CTAAGTAAGTGATATGAAGCCACATTCAAC |
| CsaCCR5-FL-For | ATGGCAACTAGCAAGGAGAAAGAGCC |
| CsaCCR5-FL-Rev | TCACTGATTCAAAAAGCCTTTGCTTCT |
| CsaCCR7-FL-For | ATGCCGATTGACGATACCTCTTCAG |
| CsaCCR7-FL-Rev | TCACCGTTGAAGTTGGGAAGGAAGT |
| CsaCCR9-FL-For | ATGCCCCTCGCTGCCC |
| CsaCCR9-FL-Rev | TTAAGCAGGAAGTGGAAGATGACCCTT |
| CsaCCR11-FL-For | ATGGGGGTTGCAGGACCCG |
| CsaCCR11-FL-Rev | TTAGGGAAATGCATTAAGAGAAAAACAAC |
| CsaCCR12-FL-For | ATGGCGATGAAGGGGAAAGTTTG |
| CsaCCR12-FL-Rev | TCACTCCAGCATGCCGGCG |
| CsaCCR13-FL-For | ATGGGCGTTATATTTCGTTCTGAAGAG |
| CsaCCR13-FL-Rev | TCAGAAATCTCTGTTTTCATTCAAACATCT |
| CsaCCR14-FL-For | ATGCCAGAGTACTGCGTAACTGGCG |
| CsaCCR14-FL-Rev | TCAAAGAAATCCTTTGTCTTGGAAACTCTT |
| CsaCCR16-FL-For | ATGGAAAGTGTGGCGAAGGGG |
| CsaCCR16-FL-Rev | CTAGAGATAACCCTTCTCTTTGCAGCTTT |
| CsaCCR17-FL-For | ATGGAGCAGAACCTTGAAACCAATG |
| CsaCCR17-FL-Rev | TTAAGGTGAAGAAAGATGGCCTTGCTC |
| CsaCCR18-FL-For | ATGCCAGTCGATACTCCTTCATCTCACT |
| CsaCCR18-FL-Rev | TCAAGATTGAATTTGAATGGAATCTGGT |
| **For quantitative reverse transcription-PCR** | |
| CsaCCR7-RT-For | ATGCCGATTGACGATACCTCTTCAG |
| CsaCCR7-RT-For | TCACCGTTGAAGTTGGGAAGGAAGT |
| CsaCCR9-RT-For | AAAAACAGTGGCAGAGCAAACA |
| CsaCCR9-RT-For | TCTACCGAAGGCAGAGGGAGTC |
| CsaCCR18-RT-For | TGTGGCTCTAGCTCATATCCTTGT |
| CsaCCR18-RT-For | GTATGCTTTCTTCCTTGGGTTCA |
| **For subcellular localization constructs** | |
| CsaCCR7-Sub-For | CAGGTCGACATGCCGATTGACGATACCTCTTCAG (*Sal* Ⅰ) |

| | |
|---|---|
| CsaCCR7-Sub-Rev | ATATCTAGATCACCGTTGAAGTTGGGAAGGAAGT (*Xba* Ⅰ) |
| CsaCCR9-Sub-For | ATAGGATCCATGCCCCTCGCTGCCCCT (*Bam*H Ⅰ) |
| CsaCCR9-Sub-Rev | ATAGAATTCTTAAGCAGGAAGTGGAAGATGACCC (*Eco*R Ⅰ) |
| CsaCCR18-Sub-For | ATAAAGCTTATGCCAGTCGATACTCCTTCATCTCACT (*Hind* III) |
| CsaCCR18-Sub-Rev | ATATCTAGAAGATTGAATTTGAATGGAATCTGGT (*Xba* Ⅰ) |
| **For recombinant protein purification** | |
| CCR7-pEasy-For | ATGCCGATTGACGATACCTCTTCAG |
| CCR7-pEasy-Rev | TCACCGTTGAAGTTGGGAAGGAAGT |
| CCR9-pEasy-For | ATGCCCCTCGCTGCCCCT |
| CCR9-pEasy-Rev | AGCAGGAAGTGGAAGATGACCC |
| CCR18-pEasy-For | ATGCCAGTCGATACTCCTTCATCTCACT |
| CCR18-pEasy-Rev | AGATTGAATTTGAATGGAATCTGGT |

**Table S2. CsaCCRs subcellular localization prediction by using Traget P and Wolf PSORT software.**

| Protein name | Transcription | Length (aa) | Isoelectric points | Predicted subcellular localization |
|---|---|---|---|---|
| **CsaCCR1** | Csa1G277460.1 | 298 | 5.46 | Cyto:0.792; Cyto:11.0 |
| **CsaCCR2** | Csa1G533570.1 | 326 | 5.92 | Cyto : 0.498; Cyto :7.0 |
| **CsaCCR3** | Csa2G441240.1 | 299 | 6.31 | SP : 0.231; Chlo:5.0; Cyto : 5.0 |
| **CsaCCR4** | Csa3G098550.1 | 345 | 5.06 | Cyto : 0.953; Cyto :5.0 |
| **CsaCCR5** | Csa3G113390.1 | 327 | 6.04 | Cyto : 0.646; Cyto :6.0 |
| **CsaCCR6** | Csa3G715370.1 | 153 | 6.78 | Cyto : 0.642; Nucl : 5.0 |
| **CsaCCR7** | Csa3G716870.1 | 330 | 6.51 | Other : 0.705; Nucl : 6.0 |
| **CsaCCR8** | Csa3G717370.1 | 345 | 7.12 | Other : 0.671; Nucl : 4.0 |
| **CsaCCR9** | Csa3G717870.1 | 324 | 7.99 | SP : 0.357; Cyto :5.0 |
| **CsaCCR10** | Csa3G717880.1 | 107 | --- | Other : 0.671; Chlo : 10.0 |
| **CsaCCR11** | Csa3G837550.1 | 385 | 6.51 | Cyto : 0.84; Chlo : 3.0, Plat : 3.0 |
| **CsaCCR12** | Csa4G664500.1 | 320 | 7.05 | SP : 0.531;Chlo : 3.0, Extr : 3.0 |
| **CsaCCR13** | Csa5G524780.1 | 364 | 7.56 | mTP : 0.6; Cyto : 6.0 |
| **CsaCCR14** | Csa5G609630.1 | 321 | 6.50 | Cyto : 0.746; Cyto : 8.0 |
| **CsaCCR15** | Csa6G077390.1 | 129 | --- | Cyto : 0.886; Extr : 6.0 |
| **CsaCCR16** | Csa6G077400.1 | 329 | 5.87 | Cyto : 0.59; Chlo : 8.0 |
| **CsaCCR17** | Csa6G116160.1 | 326 | 6.59 | Cyto : 0.704; Cyto : 7.0 |
| **CsaCCR18** | Csa7G432010.1 | 339 | 6.24 | Cyto : 0.641; Chlo : 3.0, E.R : 3.0 |

Cyto, cytoplasm; Chlo, chloroplast; Nucl, nuclear; mTP: mitochondira; E.R, endoplasmic reticulum; SP: signal peptide; Plas, plasma membrane; Extr, extracellular region.
